# *Ex vivo* infection of human skin with herpes simplex virus 1 reveals mechanical wounds as insufficient entry portals via the skin surface

**DOI:** 10.1101/2021.06.02.446703

**Authors:** Nydia De La Cruz, Maureen Möckel, Lisa Wirtz, Katharina Sunaoglu, Wolfram Malter, Max Zinser, Dagmar Knebel-Mörsdorf

## Abstract

Herpes simplex virus 1 (HSV-1) enters its human host via the skin or mucosa. The open question is how the virus invades this highly protective tissue *in vivo* to approach its receptors in the epidermis and initiate infection. Here, we performed *ex vivo* infection studies in human skin to investigate how susceptible the epidermis and dermis are to HSV-1 and whether wounding facilitates viral invasion. Upon *ex vivo* infection of complete skin, only sample edges demonstrated infected cells. After removal of the dermis, HSV-1 efficiently invaded the basal layer, and from there, gained access to suprabasal layers supporting a high susceptibility of the epidermis. In contrast, only single infected cells were detected in the papillary layer of the separated dermis. Interestingly, after wounding, nearly no infection of the epidermis was observed via the skin surface. However, if the wounding of the skin samples led to breaks through the dermis, HSV-1 infected mainly keratinocytes via the wounded dermis. The application of latex beads revealed only occasional entry via the wounded dermis, however, facilitated penetration via the wounded skin surface. Thus, we suggest that the wounded human skin surface allows particle penetration but still provides barriers that prevent HSV-1 invasion.

## Introduction

Herpes simplex virus 1 (HSV-1) is a prevalent human pathogen that can cause various infections and remains latent in their host for life. During primary infection, HSV-1 invades mucosal surfaces or abraded skin and replicates largely in the epidermis, which is followed by latent infection of sensory neurons where the virus can be reactivated to cause lesions at or near the site of initial infection. As the extent of primary and recurrent infections is largely a function of the host’s immune status, severe HSV-1 infections can occur in immunocompromised hosts and newborns. Patients with skin lesions are predisposed to primary and recurrent HSV-1 infections indicating that the virus needs to overcome skin barriers for efficient infection. After penetration of the tissue, HSV-1 entry requires the interaction of multiple viral glycoproteins with host receptors (Heldwein and Krummenacher, 2008). Upon attachment to heparan sulfate proteoglycans on the cell surface, the viral glycoprotein gD interacts with its host receptors, which in turn initiates the fusion of the viral envelope with cellular membranes (Connolly et al., 2021). The primary gD receptors for HSV-1 on human cells are the cell-cell adhesion protein nectin-1 and herpesvirus entry mediator (HVEM), a member of the tumor necrosis factor receptor superfamily (Montgomery et al., 1996; Geraghty et al., 1998). Receptor interactions are well studied in cultured cells, however, we know less about their relevance for viral entry in human skin and mucosa, and about the conditions under which the receptors are accessible *in vivo*.

In general, the epidermis as the outermost layer of the skin provides efficient barriers which prevent water loss, exclude toxins and pathogens, resist mechanical stress and participate in immune responses (Simpson et al., 2011). The initial epidermal barriers towards pathogens include chemical barriers based on antimicrobial peptides, and a physical protection based on lipid sealed cell-cell contacts in the uppermost, cornified layer. These barriers in the outer shield of the epidermis provide an outside-in barrier function. The integrity of the underlying viable epidermal layers provides a further protective physical barrier mediated by cell adhesion molecules. Key players are the tight junction (TJ) proteins which are restricted to the granular layer, the uppermost nucleated epidermal layer. TJs control the paracellular transport of molecules and thereby also form an inside-out barrier. How HSV-1 bypasses the epidermal barriers to target its receptors, either during primary or recurrent infection, is poorly understood. In order to dissect the relevance of the physical barrier functions for HSV-1 infection, we established an *ex vivo* infection model of murine skin (Petermann et al., 2009; Rahn et al., 2015a). While murine epidermal sheets are highly susceptible to HSV-1, murine total skin samples are protected against *ex vivo* infection confirming that the virus cannot penetrate via the apical skin surface (Rahn et al., 2015b). Even after removal of the cornified layer, no infected cells were observed (Rahn et al., 2017). Next to the cornified layer, functional TJs can also interfere with the efficiency of HSV-1 entry (Rahn et al., 2017).

Here, we explored the impact of physical skin barriers in human skin by *ex vivo* infection to identify the conditions under which HSV-1 can invade the tissue. Only after removal of the dermis from human skin samples and infection of the epidermal sheets was efficient infection detected in basal and suprabasal keratinocytes. Upon wounding, viral invasion was not observed via the skin surface but only when wounds crossed the dermis so that the virus gained access to keratinocytes via the dermal layer. Our results demonstrate that mechanical wounds of the skin surface do not provide *ad hoc* entry portals for HSV-1. Wounded skin samples that allowed entry via a damaged dermis and basement membrane support, however, that mechanical wounding can be sufficient for successful *ex vivo* infection with HSV-1.

## Results

### *Ex vivo* infection of human skin

To investigate the susceptibility of human skin to HSV-1, we began with *ex vivo* infection of total human skin samples by submerging them in virus suspension (Fig. 1a). We determined viral entry in individual cells by visualizing the very early expressed viral protein ICP0 to identify primary entry portals. Once the viral genome is released into the nucleus, ICP0 first localizes in the nucleus and then relocalizes to the cytoplasm during later infection, indicating viral replication (Petermann et al., 2009; Lopez et al., 2001). Thus, punctate nuclear ICP0 staining visualizes the completion of successful viral entry and cytoplasmic ICP0 indicates the onset of infection (Fig. 1b, c). Analyses of infected skin samples (n=10) revealed no ICP0-expressing cells until 16 (n=7) and 24 (n=9) hours p.i., however, only at the sample edges (Fig. 1b). Staining of collagenVII marked the basement membrane and demonstrated the loss of tissue integrity at the infected sample edges over time (Fig. 1b). After infection, the dermis was separated from the epidermis to visualize the highly susceptible keratinocytes at the edges in whole mount preparations (Fig. 1a, c). Whereas basal keratinocytes directly at the edge showed cytoplasmic ICP0 pointing to viral replication, the adjacent basal cells demonstrated nuclear ICP0 suggesting virus spreading (Fig. 1c). In contrast to the keratinocytes, we observed infection of dermal cells neither at the sample edges (Fig. 1b) nor at the bottom of the dermis. These results confirmed the efficient barrier function of the skin surface and revealed infection only in keratinocytes at the sample edges.

**Figure 1.**
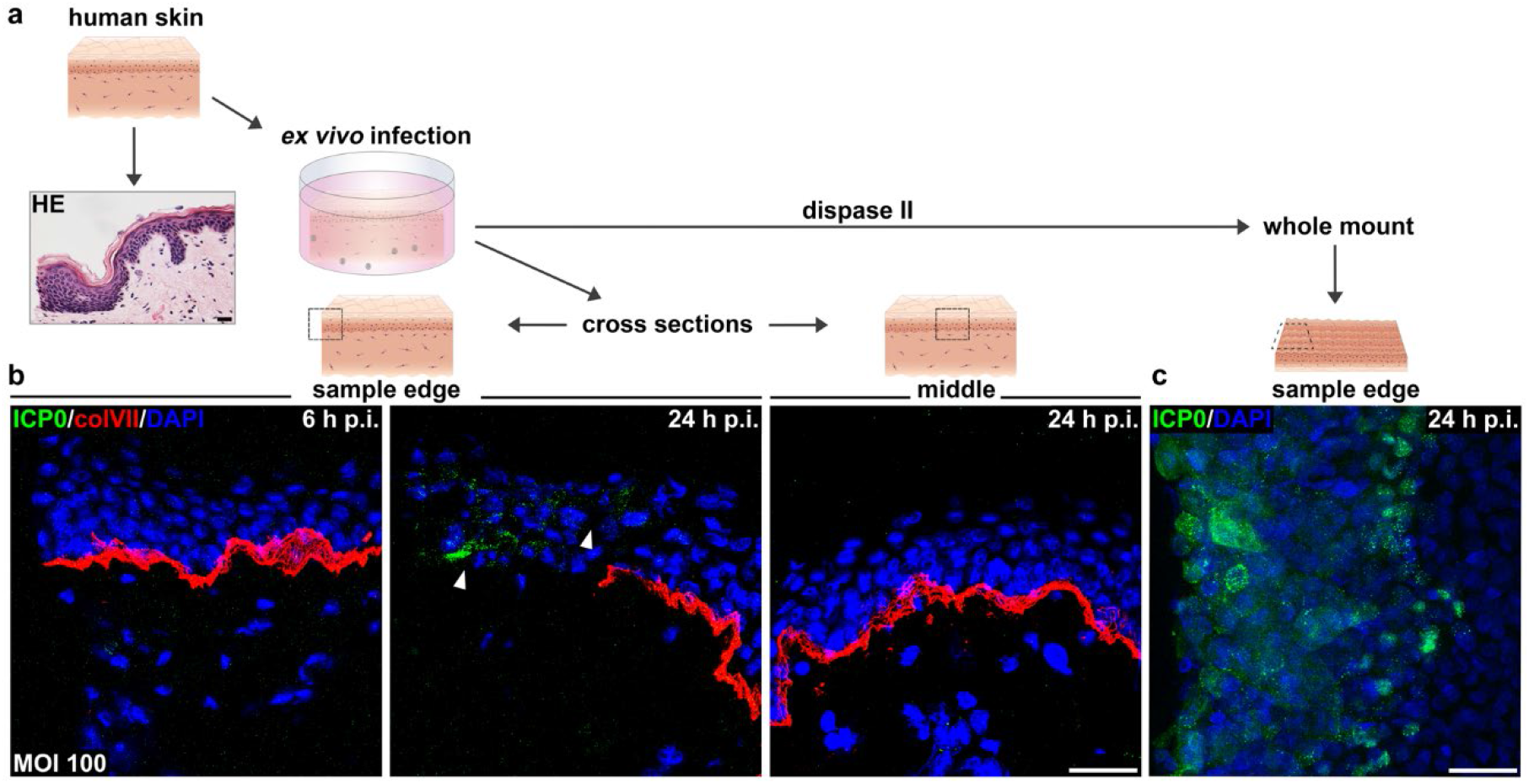
HSV-1 entry in human skin only at the sample edges. (a) Scheme illustrating *ex vivo* infection and analyses of human total skin samples. HE-stained section visualizes a sample edge prior to incubation. (b) Immunostainings of representative cross sections of breast skin (n=10) infected with 100 PFU/cell show ICP0-expressing cells (green) at sample edge only at 24 hours p.i. (arrowheads). (c) Whole mount prepared after separation of the dermis from infected total breast skin demonstrates cytoplasmic ICP0 in cells of the basal epidermal layer at sample edge and nuclear ICP0 in cells closer to the middle of the epidermis. CollagenVII (colVII) (red) depicts the basement membrane, DAPI (blue) serves as nuclear counterstain. Bars=25 μm.

### Susceptibility of human epidermal sheets to HSV-1 and viral spreading to suprabasal layers in the absence of viral replication

To address the susceptibility of the epidermis to HSV-1 in detail, we *ex vivo* infected epidermal sheets after removal of the dermis (Fig. 2a). Single cells with nuclear ICP0 were detected in the basal layer of 18 samples already at 3 hours p.i. (Fig. 2d). The number of infected cells increased at 6 hours p.i. (n=14) (Fig. 2d), and in many samples (n=12), areas with nearly all basal keratinocytes infected were observed. At 9 hours p.i., infected cells started to appear in the suprabasal layers of most samples (n=10) in addition to rather complete infection of the basal layer (Fig. 2d, e). Quantification of the ICP0 signals in the basal layer of 5 skin samples illustrates the increased susceptibility until 9 hours p.i. (Fig. 2f). As a control, stainings of cleaved caspase 3 (cc3) confirmed the viability of nearly all basal keratinocytes although in less infected areas, single cc3-positive cells were found (Fig. 2e). The distribution of infected cells in the suprabasal layers was depicted by staining loricrin as marker for terminally differentiated cells in the granular layer. The number of infected cells in the suprabasal layer increased at 12 hours p.i., while some upper granular cells expressing ICP0 were observed only at 24 hours p.i. (Fig. 2d). At least in some areas of the epidermal sheets, infection for 24 hours resulted in cytopathic effects (CPE) which may facilitate viral penetration even in the granular layer (Fig. 2d). Susceptibility of all nucleated epidermal layers to HSV-1 in human epidermis also occurs in murine epidermis (Petermann et al., 2015; Rahn et al., 2015b).

**Figure 2.**
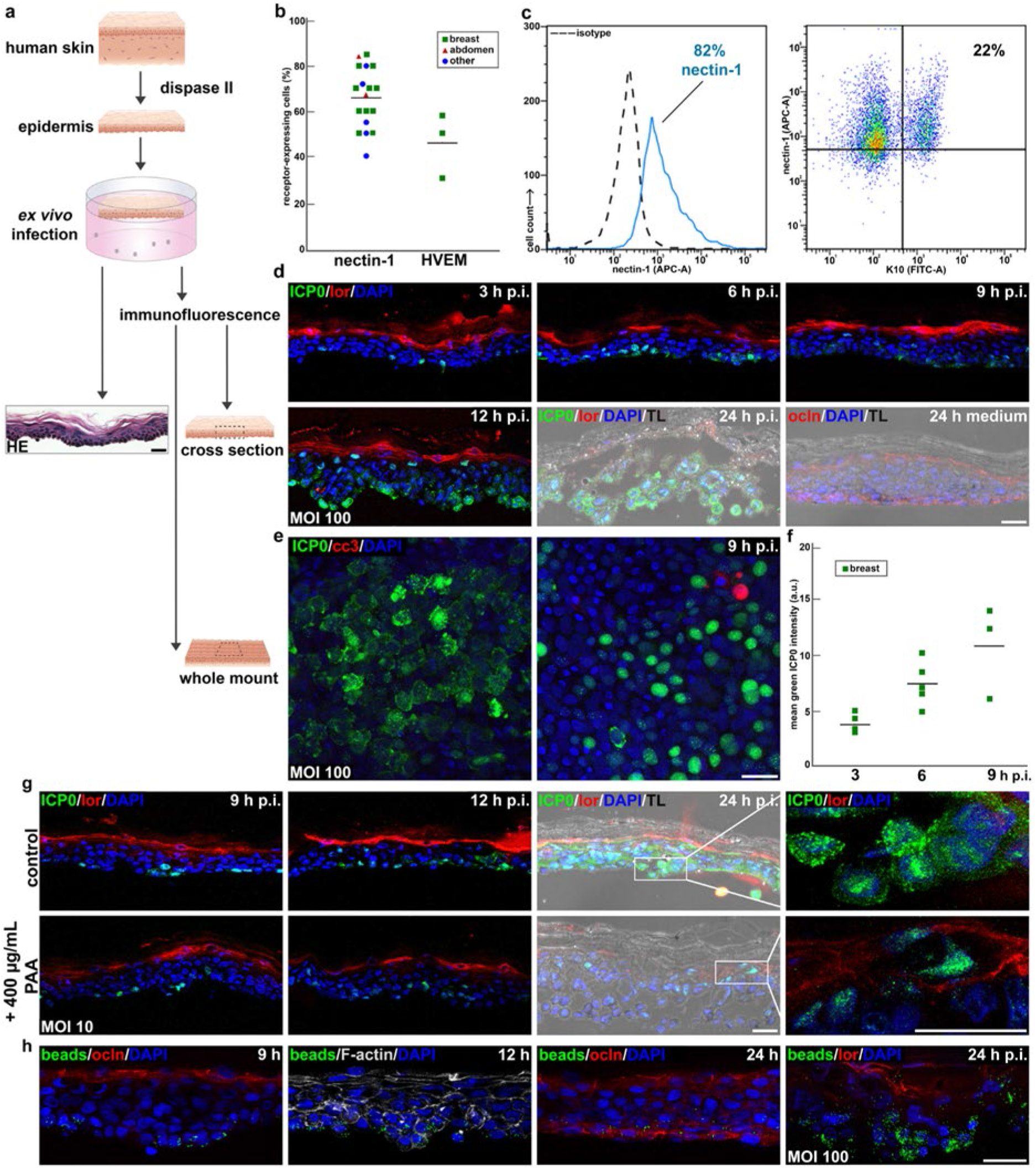
HSV-1 entry and uptake of latex beads in human epidermis. (a) Infection and analyses of epidermis. (b) Flow cytometry shows nectin-1- or HVEM-positive epidermal cells. (c) Nectin-1/K10-positive epidermal cells from breast sample. (d) Representative time course of ICP0-expressing (green) basal and suprabasal cells in abdominal epidermis (n=3); loricrin (lor) (red) depicts granular layer; occludin (ocln) (red) visualizes TJs in basal layer after 24 hours incubation. (e) Whole mount of breast skin with cytoplasmic ICP0; cleaved caspase 3 (cc3)-positive cells (red) only in area with nuclear ICP0. (f) Increased infection efficiency in breast skin over time. (g) Infection of PAA-treated abdominal epidermis results in nuclear ICP0 in basal and suprabasal cells. (h) Beads (green) uptake in suprabasal cells of abdominal epidermis after 12, no increase after 24 hours. Enhanced internalization of beads in suprabasal cells after infection. F-actin (white) depicts cell morphology. TL= transmission light. Bars=25 μm.

To distinguish between spreading of viral progeny to suprabasal cells and facilitated penetration of input virus in the epidermis during initial infection, we inhibited HSV-1 replication by phosphonoacetic acid (PAA). The successful block of viral replication is visualized by the lack of cytoplasmic translocation of ICP0 that is retained in the nucleus (Lopez et al., 2001). Upon infection of epidermal sheets for 9 hours, comparable numbers of basal cells with nuclear ICP0 were observed in the presence and absence of PAA (Fig. 2g). Interestingly, at 12 hours p.i. some nuclear ICP0-expressing cells were detected in the suprabasal layers in all samples in addition to the infected basal cells when PAA was present (Fig. 2g). At 24 hours p.i. the number of suprabasal cells with nuclear ICP0 increased and even in the granular layer, nuclear ICP0-expressing cells were found upon PAA-treatment; in the absence of PAA, all basal and suprabasal cells showed cytoplasmic ICP0 as expected (Fig. 2g). Surprisingly, we still observed some areas with disturbed morphology at 24 hours p.i. when viral replication was blocked (Fig. 2g), while tissue integrity was completely maintained after 24 hours incubation in virus-free medium (Fig. 2d). In summary, the inhibitor studies demonstrate that HSV-1 can reach suprabasal cells in the absence of viral replication at 12 hours p.i. supporting that the viral particles are able to penetrate in deeper tissue layers via the basal layer. Suprabasal ICP0 expression at 24 hours p.i. however, may be related to the tissue damage occurring not only in the presence but also in the absence of viral replication.

To explore whether virus-induced tissue permeability is involved in facilitated viral entry, we analyzed how well fluorescently-labeled latex beads (500 nm) penetrated the epidermal layers. While beads were present in basal cells after 3 and 6 hours incubation (data not shown), beads in some suprabasal cells were observed after 9 and 12 hours (Fig. 2h). However, there was no further increase of beads in suprabasal cells after 12 hours (Fig. 2h). As already observed in murine epidermis (Rahn et al., 2015b), incubation of human epidermis in medium for 24 h resulted in an extensive relocalization of TJs to the basal layer as visualized by occludin staining (Fig. 2d, h), which was not found at 24 hours after infection because of virus-induced morphology disturbance (data not shown). This remodeling of TJs was not yet visible after 9 to 12 hours thus allowing beads to be internalized in suprabasal cells only until 12 hours (Fig. 2h).

When epidermal sheets were simultaneously infected and incubated with beads, the access of beads to suprabasal including granular cells was strongly enhanced (Fig. 2h). Thus, we conclude that viral access to suprabasal layers depends largely on virus-induced tissue damage. The infection studies indicated that HSV-1 could enter all nucleated keratinocytes in the human epidermis supporting that the virus gained access to ubiquitously expressed entry receptors. Previously, we found that HSV-1 entry in murine epidermis strongly depends on nectin-1, but HVEM can potentially replace nectin-1 as receptor (Petermann et al., 2015). Thus, we investigated to which extent nectin-1 and HVEM were expressed on epidermal cells prepared from human skin, which originated from 39 to 86 year-old patients and included mainly breast (n=11) but also abdominal skin (n=2) and skin from other areas (n=4). The skin analyses by flow cytometry revealed nectin-1 on approximately 40% to 85% of the analyzed epidermal cells (Fig. 2b). The variation of nectin-1 expression correlated neither with age (data not shown) nor skin area. Co-staining of nectin-1 and keratin-10 (K10), a marker for differentiating keratinocytes, demonstrated that nectin-1 was highly expressed on basal but also differentiating keratinocytes as from 82% nectin-1-positive cells, a subpopulation of approximately 22% was nectin-1- and K10-positive indicative for suprabasal cells (Fig. 2c). This finding correlates with viral susceptibility of suprabasal cells over time. Interestingly, we observed no increased susceptibility in those epidermal sheets with very high nectin-1 expression (≥70%). In addition, the alternative receptor HVEM was expressed on approximately 35% to 60% of the basal cells suggesting that both nectin-1 and HVEM could act as entry receptors (Fig. 2b).

### Susceptibility of human dermis to HSV-1

As no infected dermal cells were observed upon infection of total human skin samples, we investigated whether the dermis is susceptible to HSV-1 after separation from the epidermis by dispase II treatment (Fig. 3a). To demonstrate whether infection led to tissue alterations, we visualized the organization of collagen fibrils by SHG microscopy which demonstrated no obvious changes of the collagen matrix after *ex vivo* infection for 24 hours (Fig. 3b). Interestingly, upon infection we detected no ICP0-expressing cells at 6 and 12 hours p.i. Only at 16 hours p.i. were single infected cells found in the most apical part of the papillary dermis with no increase at 24 hours p.i. (Fig. 3c). Co-staining of ICP0 and the leucocyte-specific surface antigen CD45 (Thomas, 1989) revealed no preferred infection of immune cells supporting that dermal fibroblasts represent the few infected cells (Fig. 3c). This rather low infection efficiency is in contrast to murine dermis, where most of the apical papillary dermis is infected at 20 hours p.i. (Wirtz et al., 2020).

**Figure 3.**
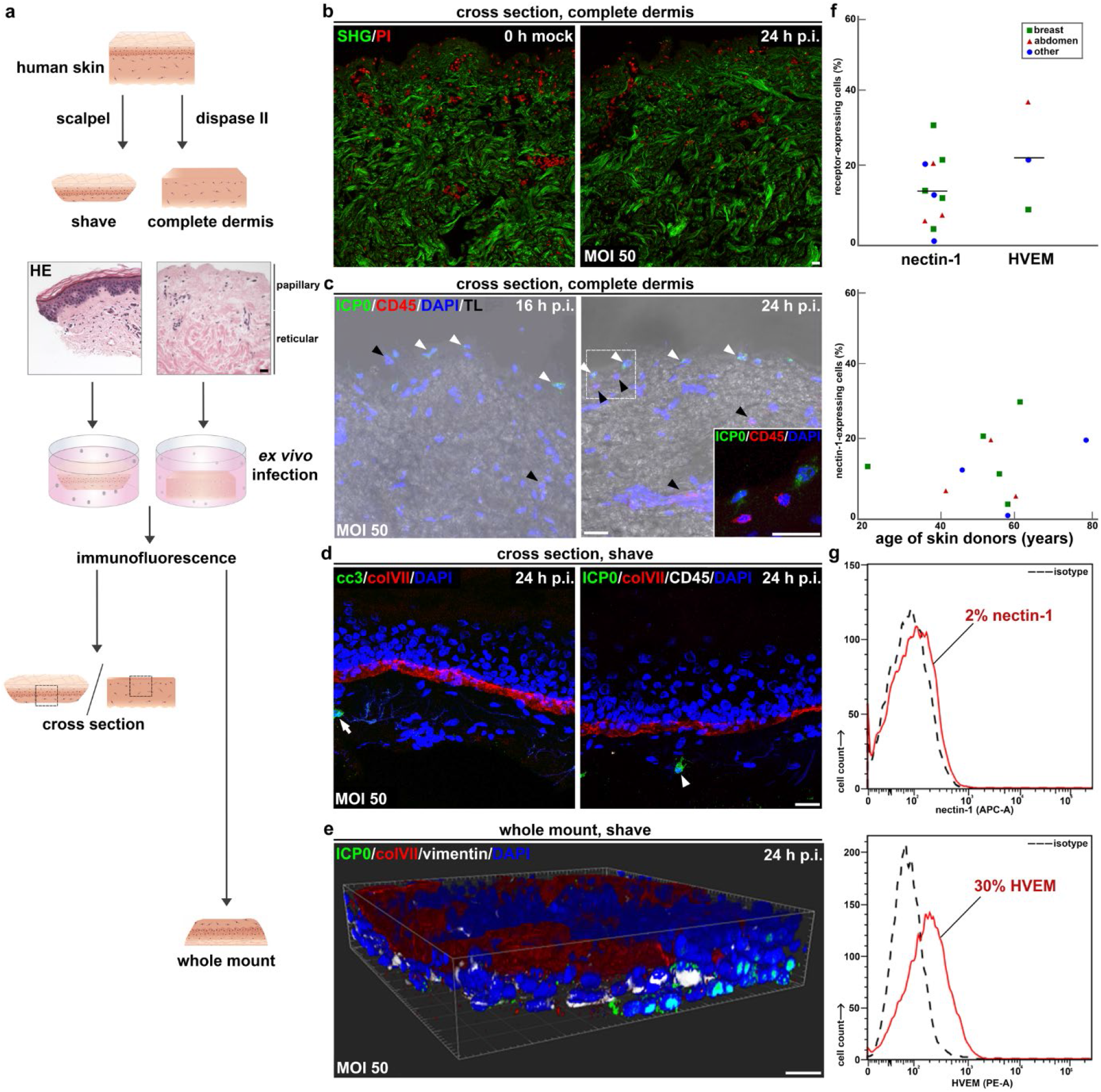
Insufficient HSV-1 entry in human dermis. (a) Infection and analyses of skin shaves and dermis. (b) Second harmonic generation (SHG) shows unchanged collagen structure 24 hours p.i. (c) Representative immunostaining of 9 samples shows single ICP0-expressing cells (white arrowheads) in the papillary layer of abdominal dermis. No costaining of ICP0- and CD45-positive (red) (black arrowheads) cells. (d) Single cc3-positive cell (arrow) and single ICP0-expressing/CD45-negative cell (arrowhead) in dermal layer of abdominal shaves. (e) 3D whole mount of infected shave visualizes vimentin-positive (white)/ICP0-expressing fibroblasts and infected basal keratinocytes at sample edge. Bars=25 μm (f) Flow cytometry demonstrates nectin-1- or HVEM-positive cells isolated from complete dermis; nectin-1 expression at different ages. (g) Nectin-1- or HVEM-positive cells from abdominal skin shave shown in d and e.

To correlate the low infection efficiency with receptor presence, we investigated surface expression of nectin-1 by flow cytometry. The analyses of dermal sheets (n=11) revealed variable nectin-1 expression ranging from undetectable to 30% of dermal cells, which correlated neither with skin area nor age (Fig. 3f). Detectable levels of nectin-1 transcripts were found in all dermal sheets (data not shown). Furthermore, we observed surface expression of HVEM on ca. 22% of dermal cells (Fig. 3f). These results suggest that the minor expression of nectin-1 and HVEM on dermal cells contributes to the low infection efficiency. To explore whether removal of the reticular dermis facilitates viral access to cells in the papillary dermis, we used skin shaves for *ex vivo* infection studies (Fig. 3a). After removal of the papillary dermis from skin shaves, we observed approximately 2% nectin-1- and 30% HVEM-expressing dermal cells (Fig. 3g). Upon infection of these shaves, single ICP0-expressing cells were found at 16 and 24 hours p.i. in the papillary dermis, which did not represent immune cells as shown by negative CD45 staining (Fig. 3d). Control staining of cc3 confirmed the viability of the dermal cells (Fig. 3d). As expected at 24 hours p.i., ICP0-expressing cells were evident at the edges of the skin shaves and mostly represented keratinocytes although co-staining of vimentin revealed also some infected dermal cells (Fig. 3e). Taken together, viral access to the papillary dermis of skin shaves was still rather inefficient.

### HSV-1 entry and latex beads uptake in wounded human skin

To address whether dysfunctional physical barriers of the skin facilitate HSV-1 entry, we applied various protocols to wound human skin samples. Immediately after wounding, the samples were infected by submerging in virus suspension or topical application (Fig. 4a). Wounding approaches such as scalpel cuts and sandpaper revealed no ICP0-expressing cells even at 24 hours p.i. (data not shown). We thus used a more reproducible method and applied microneedles, which generated lesions providing viral access to the various epidermal layers and the dermis. We *ex vivo* infected skin samples (n=14) from various areas and ages immediately after wounding. Histological sections revealed open lesions immediately after wounding (Fig. 4a), but rarely after incubation in medium suggesting that the lesions fold back after incubation. However, lesions were visible by the discontinuous basement membrane as visualized by collagenVII staining (Fig. 4b). Surprisingly, no infected cells were detected at wounded areas of 13 samples at 16 and 24 hours p.i. (Fig. 4b), but mostly only at sample edges as observed in unwounded skin (Fig. 1b). The viability of cells in the wounded area was confirmed by cc3 staining (Fig. 4b). Only in one skin sample, ICP0-expressing cells were found in a lesion (Fig. 4c) suggesting that an infected wound is a very rare event. We next infected wounded skin shaves taken from abdominal skin (n=3) (Fig. 4d). Skin shaves included the complete epidermis with a small dermal layer so that the microneedle-induced lesions completely penetrated the epidermis as well as the dermal layer. Again, we observed infected cells at the edges of most samples where tissue integrity was frequently lost at 24 hours p.i. (Fig. 4e). In contrast to total wounded skin, however, we easily detected infected cells in the wounded areas at 16 hours p.i. with increasing numbers at 24 hours p.i. (Fig. 4f). These infected cells rarely included ICP0-expressing fibroblasts, but mostly represented basal keratinocytes as visualized by whole mount preparations after removal of the dermal layer (Fig. 4g).

**Figure 4.**
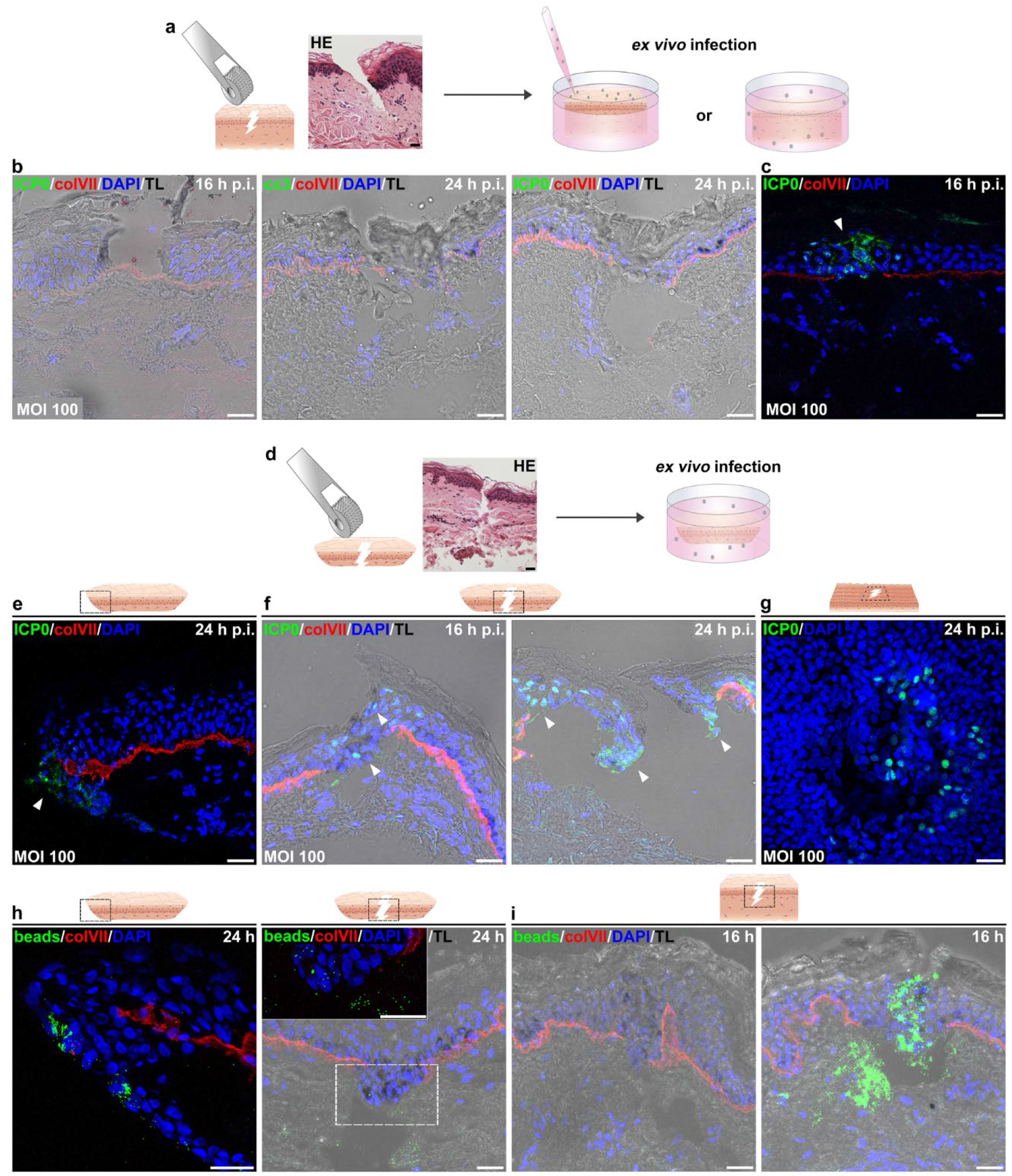
HSV-1 entry only in wounded human skin shaves but not in lesions of total skin. (a, d) Wounding, infection, and analyses of total skin and skin shaves. HE-stained sections visualize lesions prior to incubation. (b) Various lesions in total abdominal skin show neither ICP0- nor cc3-positive cells indicating non-infected, viable cells. (c) Wounded breast skin shows ICP0-expressing keratinocytes in area with discontinuous collagenVII staining. (e) ICP0-expressing cells (arrowheads) at edge, or (f) closed and partly open lesions of abdominal skin shave. (g) Whole mount after separation of dermal layer shows ICP0-expressing basal cells in wounded area of abdominal skin. (h) Internalized beads at edge and single beads in partly open lesion of abdominal skin shave. (i) No beads in closed and clustered beads in partly open lesion of total abdominal skin. Bars=25 μm

Taken together, wounding by microneedles revealed that physical lesions did not allow invasion of HSV-1 via the skin surface but when the lesions crossed the dermal layer in skin shaves, HSV-1 gained access to the epidermis. The open question is why the virus particles cannot penetrate the epidermis via the apical surface of the wounds. To address whether the lesions in total skin are insufficient to allow penetration of particles in general, we investigated whether latex beads (500 nm) can invade via lesions. Internalized beads were easily found at the edges of skin shaves after 24 hours incubation; however, only a limited number of beads were present in the wounded area of shaves (Fig. 4h). In contrast, we observed clusters of internalized beads in partly open lesions of total skin after 16 hours incubation (Fig. 4i). In more closed lesions, only identified by discontinuous collagenVII staining, nearly no beads were visible (Fig. 4i). This indicates that some lesions in total skin allow the strong penetration of beads via the wounded surface. The clustered beads might result from the retention of the particles in these lesions thus allowing enhanced uptake. In contrast, the beads can escape in lesions of shaves as they are open through the apical surface and the dermis.

In summary, we conclude that lesions in total skin provide no entry portal for HSV-1 although beads (500 nm) that are larger than virus particles (200 nm) can penetrate in cells of the wounded area. When HSV-1 can gain access to the lesions via the dermal site of skin shaves, however, infection of the wounded area is possible.

## Discussion

Viral entry is mostly well-studied with respect to virus-receptor interactions, but the missing link is to understand how the virus invades tissue to initiate infection. Here, we explored the conditions that allow HSV-1 to reach its host receptors in tissue. Our *ex vivo* infection studies in human skin samples demonstrated susceptible basal as well as suprabasal keratinocytes while entry in dermal fibroblasts was rarely detected. The high susceptibility of keratinocytes, however, was only achieved after separation of the dermis from the epidermis which allowed direct viral access to basal keratinocytes in the absence of the basement membrane. In contrast, HSV-1 infection of the dermis in the absence of the basement membrane only resulted in rare infected cells in the papillary dermis. A possible explanation could be the low surface expression of nectin-1 and HVEM on cells in the human dermis. In juvenile murine dermis, we previously observed nectin-1 on ca. 40% of the dermal cells and found more infected cells than in human dermis (Wirtz et al., 2020). During aging, however, nectin-1 expression drops while HVEM remains high in old murine dermis which correlates with delayed infection efficiency (Wirtz et al., 2020) supporting a link of receptor presence and infection efficiency. Furthermore, we speculate that the ECM provides a barrier that prevents the virus from efficiently reaching receptors on dermal fibroblasts. In human epidermis, we found a correlation of highly susceptible basal keratinocytes and high nectin-1 expression. Although nectin-1 was also present on suprabasal cells, how well the receptor is distributed in each of the suprabasal layers is open for future investigation.

To study how tissue integrity limits viral access to suprabasal layers after infection of epidermal sheets via the basal layer, we blocked viral replication to minimize virus-induced tissue damage. Intriguingly, the virus still entered granular cells over time. As some disturbed morphology also took place in the absence of viral replication, however, we assume that even early infection leads to some tissue damage including dysfunctional junctions and thereby offers access to receptors on upper granular cells. To further explore tissue permeability, we applied latex beads and observed internalized beads in suprabasal cells supporting that 500 nm particles can overcome junctions up to the upper granular layer. Simultaneous infection and incubation with beads, in turn, illustrates that virus-induced changes strongly enhance bead internalization in upper suprabasal cells. Taken together, our results suggest that HSV-1 readily accesses basal cells upon infection of epidermal sheets, then further viral invasion depends on virus-induced changes during early infection that even allow the virus to overcome the inside-out barrier formed by TJs.

Infection of total skin resulted in no viral penetration via the apical surface as expected. Infected cells were found at sample edges, which were first observed at 16 hours p.i. This was strongly delayed compared to epidermal sheets where infected basal cells were already observed at 3 hours p.i. Based on stainings of the basement membrane, we assume that incubation in medium alters tissue integrity at the sample edges over time which then allows the virus to gain access to its receptors on keratinocytes.

The general assumption is that skin lesions can serve as entry portals for HSV-1 as the cell’s status influences its plasma membrane dynamics which in turn might facilitate receptor accessibility. So far, mechanical wounding of human oral mucosa samples is insufficient for HSV-1 invasion suggesting that further contributions, such as the biofilm of the oral cavity, play a role in viral entry *in vivo* (Thier et al., 2017). Microneedle-treatment of human abdominal skin was recently described to allow productive HSV-1 infection thus providing an infection model to study antiviral compounds (Tajpara et al., 2019). The emphasis of that study was to prepare skin samples which show the pathogenesis of HSV-1 four days after infection irrespective of the initial entry portal (Tajpara et al., 2019). We also applied microneedle-treated skin, however, the question focused on whether and how HSV-1 entered cells at lesions, thus we compared wounded total skin with skin shaves. Our finding is that the wounded area of lesions in total skin shows ICP0-expressing cells only very rarely, while infected cells were detected in skin shaves where the lesion penetrated the epidermis as well as the dermis. Thus, we conclude that HSV-1 cannot enter the viable cells at lesions via the apical surface even when the basement membrane is disrupted but needs access to the basal keratinocytes via the wounded dermis. This conclusion is in line with the observations of Tajpara et al. (2019) who infected thin dermatome-cut skin submerged in medium supporting invasion via the dermal layer and the edges.

When we addressed whether the barriers restricting viral invasion via the wounded surface of total skin also hindered particle penetration, we surprisingly found clusters of internalized latex beads in some lesions but no beads in other more closed lesions. Both kinds of lesions harbored nearly no infected cells suggesting that wound closure does not interfere with viral invasion. Based on the observation that only beads but no virus particles can penetrate via the wounded surface, we assume that lesions in skin shaves and total skin provide different barrier functions to efficient viral invasion which still have to be identified. These different barriers might involve the variable accessibility and/or distribution of the receptor nectin-1 on cells in lesional skin shaves and total skin, respectively.

In summary, we found no viral entry via lesions in total skin although we demonstrated the high susceptibility of all epidermal layers and the cellular accessibility for latex beads. Sample edges only served as entry portals over time suggesting that tissue integrity loss has to precede successful viral penetration. Strikingly, the virus can enter epidermal cells once the wounds in thin skin samples allow access via the disrupted basement membrane while dermal cells showed a rather low susceptibility to HSV-1 (Fig. 5).

**Figure 5.**
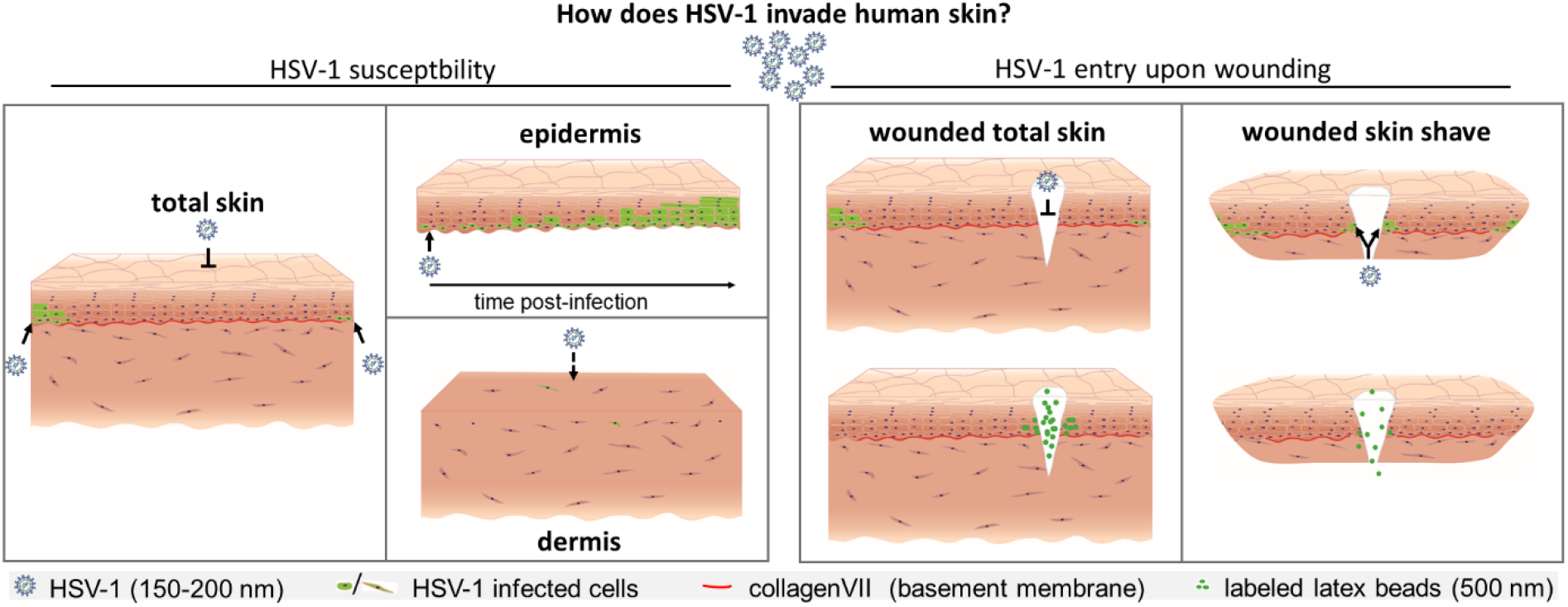
Graphical summary.

## Materials and Methods

### Preparation of human skin

Human breast (n=30) and abdominal skin (n=5) was received from patients undergoing breast and plastic surgery, respectively, while small skin biopsies (n=13) were derived from various skin areas. Immediately after surgery, skin samples with no pathological alterations were stored in DMEM/high glucose/GlutaMAX and prepared for infection. After removal of subcutaneous fat, samples were cut in pieces (total skin) or shave biopsies were taken including the epidermis and a thin dermal layer with an average thickness of 0.4 mm (skin shaves). Total skin and skin shaves were treated with a derma-roller/microneedles (1.5 mm length x 0.25 mm in diameter) (Lurrose) 5 times in 4 directions. Epidermal and dermal sheets were prepared after incubation overnight at 4°C with 4 U/ml dispase II (Roche) in PBS by gently separating dermis and epidermis each as intact sheets (Rahn et al., 2015a; Wirtz et al., 2020). All skin samples were incubated in DMEM/high glucose/GlutaMAX (Life Technologies) containing 10% FCS, penicillin (100 U/ml) and streptomycin (100 μg/ml). The analysis of viral susceptibitility revealed no detectable difference among the various skin samples.

### Ethics statement

Human skin specimens were obtained after informed consent from all patients. The study was approved by the Ethics Commission of the Medical Faculty, University of Cologne (approval no. 17-481).

### Virus

Infection was performed with purified preparations of HSV-1 wt strain 17 as described (Schelhaas et al., 2003). The calculation of the virus dose was based on the estimated cell number in the basal layer of the epidermis (ca. 3×10^5^ cells per 5- by 5-mm sheet) and in the superficial areas of the dermis (ca. 2×10^5^ cells per 4- by 4-mm surface area). Intact or microneedle-pretreated total skin and skin shaves were infected at 100 PFU/cell, dermal sheets at 50 PFU/cell, and epidermal sheets at 100 or 10 PFU/cell. The virus inoculum was added to the tissue samples at 37°C defining time point zero. Epidermal sheets were floating on virus-containing medium, while dermal sheets, total skin and skin shaves were submerged.

Viral replication was inhibited by adding PAA diluted in medium to a final concentration of 400 μg/ml (Lopez et al., 2001). PAA was present throughout infection and H2O served as the solvent control.

### Marker uptake

Sulfate-modified polystyrene, fluorescent orange latex beads (500 nm) (Sigma) served as a marker for penetration of particles in various skin samples. Total skin, skin shaves and epidermal sheets prepared from breast or abdominal skin were incubated with latex beads (2×10^9^ beads/sample) for various times at 37°C. Samples were thoroughly washed three times and immediately embedded for preparation of cryosections.

### Histochemistry, immuncytochemistry and antibodies

For cryosections, total skin, skin shaves, and epidermal or dermal sheets were embedded in OCT compound (Sakura), frozen, and cut into 8 μm (total skin, shaves, epidermal sheets) or 100 μm (dermal sheets) cross sections as described (Rahn et al., 2015a; Wirtz et al., 2020). For hematoxylin and eosin (HE) staining, samples were fixed with 3.4% formaldehyde overnight at 4°C, prepared as paraffin sections (8 μm), and stained for 10 min with hemalum followed by counterstaining with eosin for 20 seconds. HE stainings were used to assess the morphology of uninfected or infected tissue as well as of intact and microneedle-pretreated total skin/skin shaves.

For immunofluorescence, tissue sections of total skin, shaves and epidermis were fixed with 1% formaldehyde for 10 min at RT, dermal sections and whole mount preparations of shaves and epidermis were fixed with 3.4 % formaldehyde overnight at 4°C and blocked as described (Rahn et al., 2015a; Wirtz et al., 2020). Only for occludin staining, epidermal sections were fixed with 4°C cold ethanol for 30 min and then with acetone (−20°C) for 3 min. Tissue sections of total skin, shaves and epidermis were incubated with primary antibodies overnight at 4°C followed by incubation with the species-specific AlexaFluor-conjugated secondary antibodies and DAPI for 45 min at RT. Epidermal whole mounts were incubated with primary antibodies overnight at RT and with the secondary antibodies and DAPI overnight at 4°C. Dermal sections and whole mounts of shaves were incubated with primary antibodies overnight at RT and with secondary antibodies and DAPI overnight at RT. The following primary antibodies were used: mouse anti-ICP0 (monoclonal antibody 11060; 1:60) (Everett et al., 1993), mouse anti-collagenVII (1:500) (Santa Cruz Biotechnology), rabbit anti-cleaved caspase 3 (1:400) (Cell Signaling), rabbit anti-loricrin (1:1000) (Biolegend), rabbit anti-vimentin (1:400) (Cell Signaling), and mouse anti-CD45-CoraLite 488 (1:400) (Proteintech). CollagenVII staining of dermal and epidermal sections suggests the loss of the basement membrane after separation by dispase II treatment (data not shown). F-actin staining of epidermal sections was performed with phalloidin-Atto 565 (1:2000) (Sigma) for 45 min at RT to demonstrate the internalization of beads.

Second harmonic generation (SHG) microscopy was performed to analyze dermal collagen morphology by using an upright multiphoton microscope (TCS SP8 MP; Leica Microsystems) equipped with a Ti:Sa laser (Chameleon Vision II; Coherent), which was tuned to 1050 nm, as described previously (Do et al., 2018). Paraffin tissue sections co-stained with propidium iodide (PI) were placed on a mirror to improve forward directed signal detection. A 25x water immersion objective (NA 0.95, Leica Microsystems) was used and the two signals were recorded simultaneously by two non-descanned HyD detectors using a 525/50 bandpass filter for the SHG signal and a 585/40 bandpass filter for the fluorescent PI signal. LAS X software (Leica Microsystems) was used for laser scanning control and image acquisition.

Microscopy of tissue sections and whole mounts was performed using a Leica DM IRB/E microscope linked to a Leica TCS-SP/5 confocal unit. Images were assembled using Photoshop (Elements 2018; Adobe) and Illustrator (version CS2; Adobe). Images were analyzed using FiJi (version 2.0.0-rc-65/1.51s) (Schindelin et al., 2012) by measuring the mean green fluorescent intensity of three different areas per biological replicate. 3D visualization of a shave whole mount was generated with ImarisViewer (version 9.6.0, Imaris Bitplane) based on confocal z-stack acquisitions.

### Flow cytometric analysis

Epidermal and dermal sheets were prepared from human breast and abdominal skin in addition to skin from other areas. Epidermal sheets were incubated with TrypLE Select (Life Technologies) or with enzyme-free cell dissociation solution (CDS) (Sigma), and processed as described (Petermann et al., 2015). HVEM expression on cells was detected only when cells were dissociated using enzyme-free CDS. After dissociation of the epidermal sheets by enzyme-free CDS, HE stainings of the remaining sheets demonstrated that the basal cells were detached indicating that HVEM expression could only be analyzed on basal cells. In contrast, dissociation of the epidermal sheets by TrypLE Select resulted in the dissociation of basal as well as suprabasal cells although to a slightly varying extent in the different samples (data not shown). Cell suspensions prepared by TrypLE Select were incubated in PBS-5% FCS on ice for 30 min with mouse anti-nectin-1 (CK41; 1:100) (Krummenacher et al., 2000), and nectin-1 was visualized with anti-mouse IgG-Cy5 (1:100) (Jackson Immunoresearch Laboratories Inc.). After permeabilization with 0.2 % saponin, the cells were incubated in PBS-5% FCS on ice for 30 min with rabbit anti-K10 (1:1000) (Biolegend), and K10 was visualized with anti-rabbit AF488 (Life Technologies; 1:200). Cell suspensions prepared by enzyme-free CDS were kept in PBS-5% FCS and incubated on ice for 30 min with rabbit polyclonal anti-human HVEM (R140; 1:500) (Terry-Allison et al., 1998) followed by visualizing HVEM with anti-rabbit AF488 (Life Technologies; 1:200). The size of the epidermal samples mostly allowed only the analysis of nectin-1 expression. For nectin-1, mouse IgG1 (Life Technologies, 1:20) and for HVEM, rabbit IgG, polyclonal (Abcam, 1:20) were used as isotype controls.

Dermal samples and dermal sheets prepared from skin shaves were digested with whole skin dissociation kit (Miltenyi) for 2.5 hours shaking (180 rpm) at 37°C before filtering through 40 μm cell strainers. Cell suspensions were incubated in PBS-5% FCS on ice for 30 min with mouse anti-nectin-1 (CK41; 1:100) (Krummenacher et al., 2000), or with anti-HVEM-PE (Miltenyi, CD270-PE, REA247; 1:11) for 10 min at 4°C. Nectin-1 was visualized with anti mouse IgG-Cy5 (Jackson Immunoresearch Laboratories Inc.; 1:100). Because of the limited cell number prepared from dermal samples, only nectin-1 expression was analyzed in most samples. For nectin-1, mouse IgG1 (Life Technologies; 1:20) and for HVEM, PE-labelled recombinant human IgG1, (Miltenyi; REA293-PE; 1:50) were used as isotype controls. Staining with DAPI for 1 min prior to analysis allowed the gating of only viable cells. Samples were analyzed by using a FACSCanto II flow cytometer and FACSDiva (version 6.1.3, BD) and FlowJo (version 7.6.3, Tree Star) software.

## Acknowledgments

We thank Roger Everett for ICP0 antibodies, Claude Krummenacher for nectin-1 antibodies (CK41), Matthias Rübsam and Sara Wickström for critical reading of the manuscript, Christian Jüngst/CECAD Imaging facility for help with SHG microscopy. Additional thanks go to Just Vlak and Adelheid Elbe-Bürger for discussion.

This research was supported by the Deutsche Forschungsgemeinschaft (SFB829 Z4 project, KN536/16-3 to DKM, SFB829 stipend to NDLC), the Köln Fortune Program/Faculty of Medicine, University of Cologne, and the Maria-Pesch foundation.

